# Modeling the phage properties best for therapy

**DOI:** 10.64898/2025.12.21.695819

**Authors:** James J Bull, Gurneet Kaur, Stephen M Krone

## Abstract

The phages used to treat bacterial infections in phage therapy are commonly chosen based on their abilities to form plaques on the infecting bacterium – on host range. In practice, phage therapy is not always successful, leaving room for improvement. Here we use computational models to investigate whether some standard phage properties (burst size, lysis, adsorption, decay rate, growth rate) might serve as predictors of treatment success. As our measure of treatment success, we deviate from many other approaches by calculating the number of phage needed to suppress bacterial densities 100-fold in the short term, given that the patient’s immune system is expected to regain control once bacterial numbers are reduced. Numerical analysis of 2400 combinations of different values of phage phenotypes reveals that, on average, adsorption rate and growth rate provide the most useful predictive values, decay rate provides some value, whereas burst size and lysis time offer essentially little or no value. Bacterial density is especially informative of the number of phage required for treatment. There is nonetheless often considerable variation around average behavior for a single phenotype. These results raise the possibility that adsorption rate and growth rate may be especially important in phage therapy performance for both high and low bacterial densities. Given that therapeutic phages are often evolved in vitro for broad host ranges rather than for individual hosts, it should be considered that selection for broad host range may have a downside of compromising adsorption to and growth rate on individual bacterial hosts.

---

The first step in treating a bacterial infection with phages is to find one or more phages that grow on the infecting bacterium, which is not always trivial [35, 1, 16, 17, 18, 20, 34, 15]. Beyond this obvious consideration of host range, there are other attributes of the applied phages that may improve outcomes: lytic instead of temperate lifestyles [32, 15], depolymerases that degrade bacterial capsules [41, 8], slow in vivo decay rate [30, 37], and in vitro adaptation to the bacterium prior to treatment [4]. The effect of these secondary attributes has rarely been evaluated in applications, however (although the downside of using a single temperate phage is easy to anticipate a priori). Nor is it clear how one might use specific trials to evaluate the utility of these secondary attributes as generalities, as two phages that differ in one attribute might well differ in others that could confound interpretation. When considering applications of phage therapy using many different phages, how consistently beneficial is one attribute likely to be?

Here we use computational models to evaluate the consistency of various phage properties in suppressing bacterial populations. Our approach is broadly similar to that of [3], who modeled various measures of phage-bacterial growth in simulated microplate wells. We deviate from their analysis in representing a different spectrum of treatment conditions, applying a different metric of success, and incorporating an additional phage property that might be relevant in actual treatment. The comparison of both approaches should thus contribute to evaluating the robustness of these kinds of studies.

The growth of a lytic phage on bacteria depends on bacterial density and four phage characteristics: burst size, time to lysis, adsorption rate to its host, and phage death or decay rate [2, 13]. Three of these phage may change with the state of the bacteria [19, 3], but we neglect those changes here. To capture the diversity that is likely to be encountered when using a variety of wild phages, phages are here assigned any of several different burst sizes, latent periods, adsorption rates, and decay rates, each chosen from a set of values spanning a biologically-motivated range. In any trial, a single phage is assigned a single set of these four values. Collectively, trials are run for all 2400 possible combinations of the four sets of parameter values. The 2400 necessarily reflects the limited numbers of values we employ for each of the phenotypes, but it’s the span of values that should matter for our purposes rather than the density of coverage across the span. Once a phage is parameterized, it is evaluated for its ability to suppress a bacterial population by a criterion explained below. The benefit of the modeling approach is that many different phage properties can easily be compared. The models obviously differ from actual treatment in many ways, but they can nonetheless give insights that may help guide and limit empirical tests that are possible only on a much smaller scale.

## Methods

The approach here involves three conceptual steps. The first is to construct a computational model to represent an infection and possible treatments. Second, a criterion is chosen to represent success versus failure from treatment. Third, different possible treatment designs or variables (i.e., phage characteristics) are numerically investigated to decide how often success results. To apply the results, a fourth, empirical step is needed: empirically measuring the characteristics to decide which phages are best for treatment.

### The model

The computational model uses ordinary differential equations of bacterial and phage densities over time. As phage therapy is mostly applied to chronic infections [1], the rate at which bacterial densities increase will be slow compared to the span of treatment. Furthermore, bacterial densities may differ by orders of magnitude across different chronic infections. Thus, to allow for different, relatively stable bacterial densities during treatment of a chronic infection, the models assume a bacterial doubling time of 2 days; to represent different infection densities, trials will merely vary the initial bacterial density. The basic set of equations is:

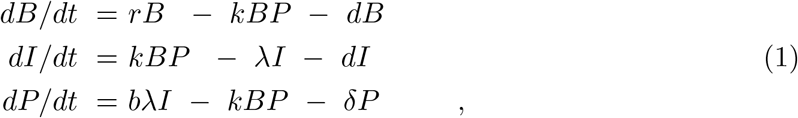

with notation and units given in Table 1. These equations are simple modifications of those used previously [33, 44]; they differ from those of many others in using a lysis rate or a gamma function instead of a fixed latent period (time from infection to lysis) [13, 27, 9, 6, 19, 3] . The ramifications of a lysis rate have been found to be largely inconsequential for our purposes [6], and a lysis rate affords considerable computational simplicity. We also note that, in contrast to the models of [19] and [3], ours neglects free phage loss to infection of already-infected cells. We have run our model to include ‘superinfection’ and found that it makes almost no difference.

**Table 1:** Model variables and parameters.

| Notation | Description | Units | Values |
| --- | --- | --- | --- |
| Variables* |  |  |  |
| $B$ | density of bacteria | /mL | $10^4, 10^6$ |
| $I$ | density of infected bacteria | /mL | |
| $P$ | density of free phage | /mL | As needed |
| Parameters |  |  |  |
| $k$ | adsorption rate constant | mL/min | $10^{-8} - 10^{-12}$ |
| $\delta$ | decay rate of free phage | /min | .001 – .01 |
| $d$ | bacterial death rate (infected and not) | /min | .001 – .01 |
| $b$ | burst size of infected cell | | 5-85 |
| $\lambda$ | lysis rate of infected cell | /min | 0.01-0.1 |
| $r$ | intrinsic bacterial growth rate | /min | 0.000241 |

\*Variables are functions of time.

Parameter ranges are chosen with long-standing empirical guidance [2, 13]. Adsorption rate has a maximum based on particle size [2] and can otherwise go to 0, although we have chosen a lower limit based on the ability to detect infection by plating [27]. Burst size can be considerably larger than we allow, but our use of a lysis rate inflates the effect of burst size over that with use of a fixed latent period [6], so we avoid burst sizes in the hundreds. Lysis rates considered here correspond to mean lysis times of 10 to 100 minutes, suitable for laboratory estimates. Decay rates have been estimated in [7] from graphs in published empirical studies [30, 45].

Aside from bacterial density, phage growth in this system is fully characterized by four phage phenotypes: burst size (*b*), adsorption rate (*k*), lysis rate (*λ*) and decay rate (*δ*). The approach used here is to assign biologically appropriate values to the four phage phenotypes (ranges given in Table 1), choose an initial bacterial density for the infection, and then numerically find the minimum phage dose added at time 0 to suppress the bacterial population 100-fold below its initial value within 2000 minutes (1.4 days). This calculation is repeated for 2400 different combinations of phage parameters and for each of two initial bacterial densities. The final data set is then analyzed for the relationship between a phage property and the minimum phage dose over the entire data set at each of the two initial bacterial densities, 10^4^ and 10^6^. This approach may be expanded indefinitely to include other bacterial densities and other values of phage phenotypes, but the sets considered here appear to provide considerable insight. Indeed, two recent modeling papers allow for considerably higher bacterial densities (10^8^ or more), as densities relevant to microplates, but infection densities may often be much lower [36, 22].

### A measure of treatment success

Whereas it is common to measure phage ‘success’ as suppression of bacteria in the long term, at equilibrium [27, 25, 26], it is the short term density that is often of interest in phage therapy – whether the phage can quickly suppress bacterial densities so that the immune response can regain control. Furthermore, bacterial densities of an infection may not be high enough to sustain the phage [33], in which case there would be no long term suppression without ongoing phage administration. In cases of low bacterial density, it is necessary to treat by administering a large dose of phage that has the short term effect of suppressing the bacterial population even though the phage will eventually die out; the same is true if applying a non-replicating phage [33, 5]. Our modeling approach searches for the minimum phage inoculum (denoted *P*_0_) needed to quickly suppress the bacterial population to 0.01 of its initial density. If the phage can amplify on the bacterial population, then administration of a single phage may suffice, but the minimum needed may otherwise be 10^7^ phage or more [e.g., 15]. Choosing success as suppression by 100-fold is necessarily somewhat arbitrary, but it is likely representative of other reasonable thresholds.

### Treatment variables

Several phage properties are evaluated here as possible determinants of phage therapy success. Four straightforward ones are the phage phenotypes of burst size, adsorption rate, lysis rate, and decay rate. The first three are easily measured under standard laboratory conditions in liquid media [2], albeit that laboratory conditions may not translate well to in vivo conditions. Decay rate is specific to a live (animal) host, although there are suggestions that it may be assessable in vitro [37]. Another possible indicator of phage success is a composite of those phage phenotypes: phage growth rate, measured as the intrinsic rate of increase. This growth rate is a measure of how fast the phage population increases when at low density [5], although it is undoubtedly highly correlated with other growth rate measures that have been used recently [19, 3]. It is highly dependent on bacterial density, so its only practical predictive value is if calculated in the lab for a standard protocol [9]. With *B* constant (set to value *C* to distinguish from the variable *B*), eqn set (1) is a linear system of two equations, and phage growth rate is given by the dominant eigenvalue:

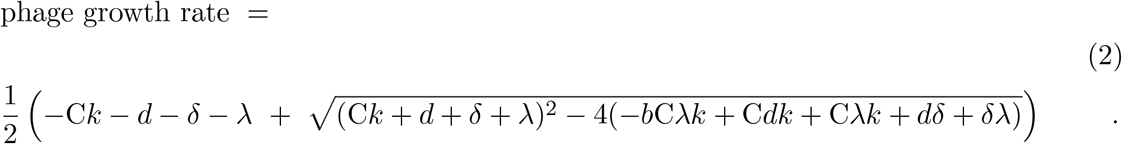

This growth rate differs from the calculation in previous studies of ours [5, 9] because the one here uses a lysis rate instead of a latent period of fixed duration. Since growth rate is feasible as a lab-determined measure of success only if calculated under a common protocol, we assume *C* = 10^8^ as a practical bacterial density to use in the lab [9], realizing that the growth rate obtained would not match growth rates for other bacterial densities that might be found in a patient. But the goal is to determine whether a growth rate obtained under one set of conditions will inform treatment success under various conditions.

Other composite measures of phage performance might also be entertained. One is the equilibrium density of cells that would occur under phage control [27]. A chemostat might be used for such a measure, but bacterial wall growth, bacterial resistance, and oscillations are potential challenges in applying this measure, even though it is easily calculated mathematically. Likewise, various measures of growth rate have been proposed for assays done in microplates using optical density of bacteria as the phenotype directly measured [19, 3].

### A second model

A second set of equations is also studied [eqn (3)]. This model differs from eqn set (1) only in that phage are introduced into a bacteria-free compartment from which they migrate to (and from) the bacterial compartment. This model is intended to mimic treatment of a patient in which phage are administered at a site remote to the bacterial infection such that there may be a delay in phage reaching their hosts:

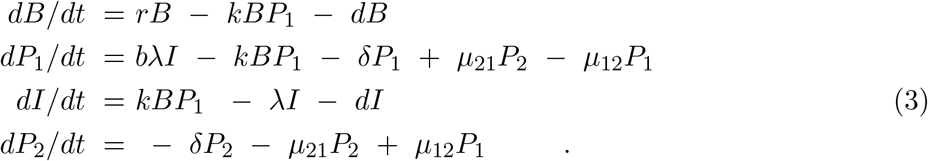

Subscripts indicate patch number of the phage (1, 2), and *µ*_*ij*_ is the migration rate from patch *i* to *j*. All bacteria, whether infected or not, are confined to patch 1.

## Results

The goal is to discover which phage properties show the strongest correlations with treatment success and, equally, to translate phage ‘quality’ into some measure of treatment effort required. With enough effort, both good and poor phages should be able to suppress an infection; the poor phage merely requires a larger inoculum, more time, or additional doses. It is treatment effort that enables us to understand how important – or not – it is to identify the best phages.

The first analyses use model (1), with results presented as plots of a single phage characteristic versus the minimum phage inoculum needed to suppress bacterial densities 100-fold (denoted *P*_0_, Figs. 1,2). Each figure has separate panels for the 5 phage characteristics considered, with the first 5-panel figure for cells at 10^6^/mL, the second figure for cells at 10^4^/mL. Each panel shows the distribution of (weighted) individual points as well as the means (rhomboids) at each of the parameter values analyzed.

**Figure 1.**
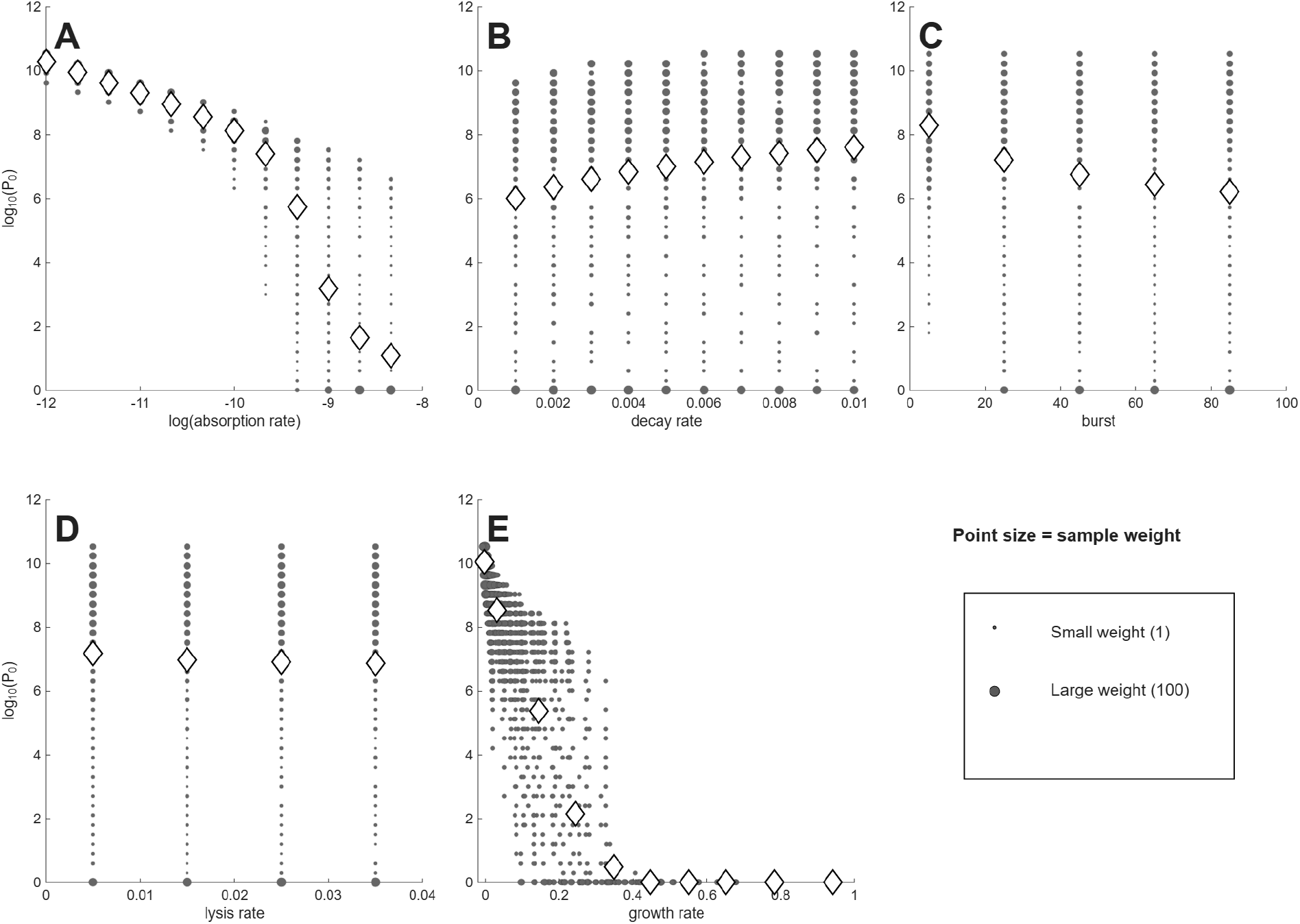
Different phage properties have different utilities as indicators of phage therapy feasibility when initial bacterial density is 10^6^; this figure is similar to that of Fig. 1, the only difference in run conditions being in the initial bacterial density. Shown are (the log of) the numbers of phage required to suppress bacterial density 100-fold (Y axis) per different phage characteristics (X axis). (A) is for adsorption rate (*k* in the model). (B) is for decay rate (*δ*). (C) is for burst (*b*). (D) is for lysis rate (*λ*). (E) is for growth rate, assuming a bacterial density of 10^8^, as per eqn (2). Only if the means span a wide range of values on the Y axis is the phenotype a good predictor of success. Thus, adsorption is best, growth rate next best. For (A) - (D), the phenotype values evaluated were each chosen at regular intervals to span a biologically-motivated interval. For (E), the growth rate values are consequences of the four phage parameter values and the 10^8^ cell density and so do not obey regular intervals. Gray filled circles represent individual values, with point diameter given the the square root of the number of data at that coordinate. Rhomboids are the means of all 2400 values numerically trialed, which often differ from the visual impression from the gray circles because the gray circles can represent many different trials.

**Figure 2.**
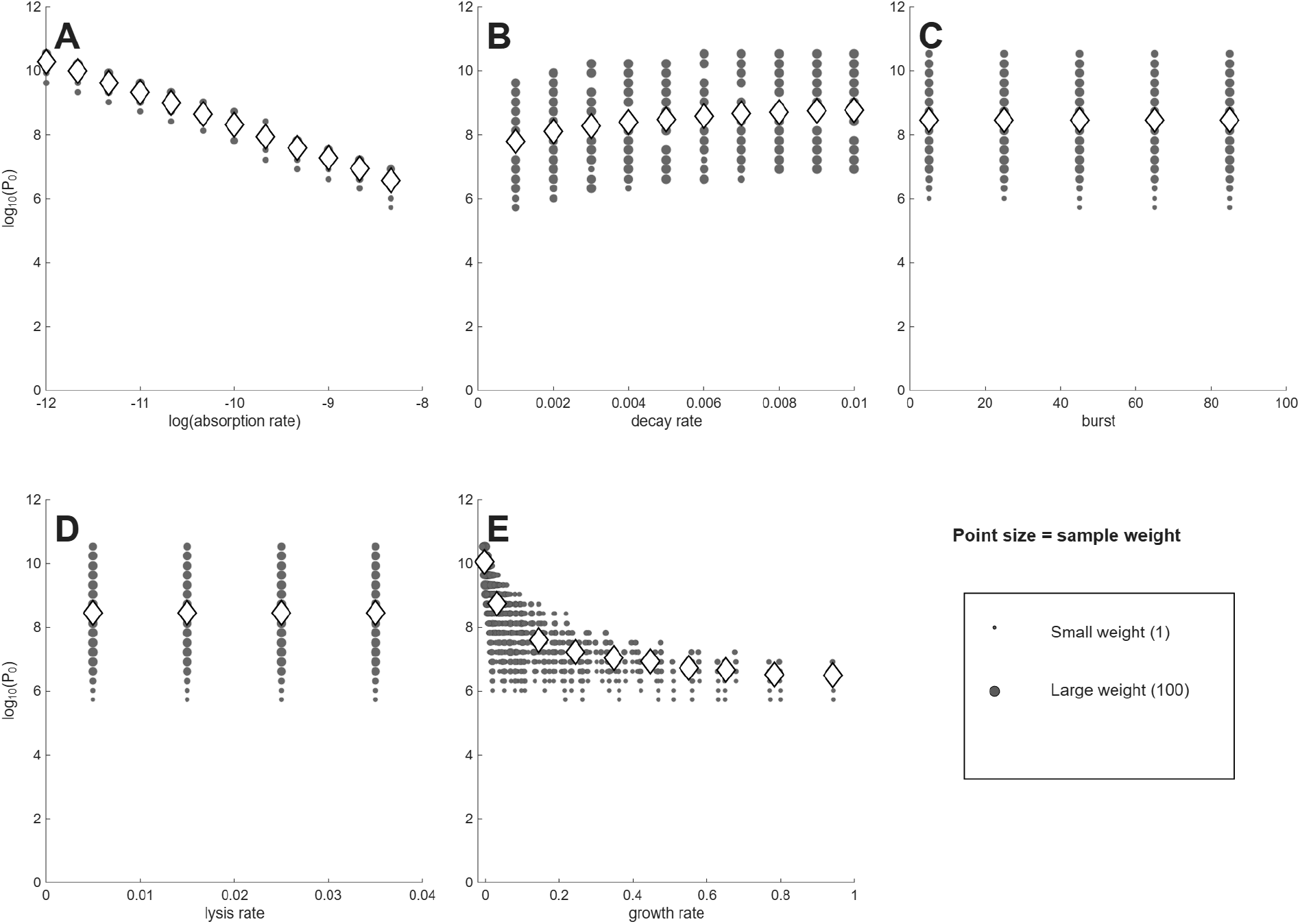
Different phage properties have different utilities as indicators of phage therapy feasibility when initial bacterial density is 10^4^. Shown are (the log of) the numbers of phage required to suppress bacterial density 100-fold (Y axis) per different phage characteristics (X axis). (A) is for adsorption rate (*k* in the model). (B) is for decay rate (*δ*). (C) is for burst (*b*). (D) is for lysis rate (*λ*). (E) is for growth rate, assuming a bacterial density of 10^8^, as per eqn (2). Only if the means span a wide range of values on the Y axis is the phenotype a good predictor of success. Thus, adsorption is best, growth rate next best. For (A) - (D), the phenotype values evaluated were each chosen at regular intervals to span a biologically-motivated interval. For (E), the growth rate values are consequences of the four phage parameter values and the 10^8^ cell density and so do not obey regular intervals. Gray filled circles represent individual values, with point diameter given the square root of the number of data at that coordinate. Rhomboids are the means of all 2400 values numerically trialed, which often differ from the visual impression from the gray circles because the gray circles can represent many different trials.

As an approximation, a log(*P*_0_) of 0-3 corresponds to ‘active’ phage therapy, in which phage amplification from a small inoculum overwhelms the bacteria, whereas high values (log(*P*_0_)> 7) correspond to ‘passive’ phage therapy, in which most cells are killed chiefly by the inoculum itself [33]. Necessarily, a combination of active and passive processes are involved for the many outcomes whose *P*_0_ values lie between. It should be noted that the set of *P*_0_ values is the same between each panel of the same figure, so the different panels merely represent different ways of sorting the same data. Likewise, the same suites of phage properties are evaluated in Fig. 1 as in Fig. 2, so differences between the two figures are due entirely to cell density. The goal is to find a way to sort the data to easily identify phages that enable active therapy, as those phages don’t require large inocula for successful treatment.

Some general points are evident from casual inspection. (1) There is no active therapy for cells at 10^4^ – no *P*_0_ values are even as low as 10^5^. In contrast, many phage combinations exhibit active therapy for cells at 10^6^, but only for the traits of adsorption rate and growth rate do the means (rhomboids) enter the realms of active therapy. Even when the means remain high for a phage characteristic (e.g., decay rate, burst size, lysis rate), individual points often range into the active therapy zone. Considering even higher cell densities (as applies in the two studies of microplate dynamics [19, 3]) would move ever more of the phages into the active realm. These comparisons highlight the necessity of matching treatment to infection conditions. (2) In comparing panels for the same phage property between the two cell densities, the *P*_0_ means for cells at 10^4^ only sometimes show the expected shift above those for 10^6^. For example, at the lowest growth rate or the lowest adsorption rate, the mean *P*_0_ is near 10 for both cell densities. For others, there is a consistent offset (e.g., lysis rate). This lack of consistency makes it difficult to develop intuition.

Not all phage properties are equal in their impact on *P*_0_. A phage property is more diagnostic of success the more that the average *P*_0_ varies across the set of phages available for treatment. It would be the distribution of values among phages obtained in nature that matters, and that distribution would likely not be uniform across the values considered here. Nonetheless, we use the span shown on the X-axis as the representative distribution, spans that were at least empirically motivated.

Of the five phage properties considered, two exhibit similar, large magnitudes of variation in average *P*_0_ across the X-axis: adsorption rate (*k*) and growth rate. For cells at 10^4^, the averages of each of those two properties spans 3.5-3.7 in log(*P*_0_) across the values given on the X-axis, whereas averages of the other three properties span no more than 1: 1.0 for decay rate, and 0 for burst and lysis. Despite the similar spans for adsorption rate and growth rate at 10^4^, the pattern for adsorption rate shows a virtually linear decline across the X-axis range, whereas the pattern for growth rate exhibits an early decline and then is flat for most of the graph. For cells at 10^6^, the spans of log(*P*_0_) are 9.2-10 for adsorption and growth rate but are 1.6, 2.0 and 0.3 for decay, burst, and lysis. Because burst and lysis time have such limited effect on *P*_0_, they will not be discussed further. We note, again, that there is considerable scatter, and the average *P*_0_ values do not necessarily represent most of the phages at a given phenotype.

A span of 9-10 in log(*P*_0_) occurs across the spectrum for two of the averages with cells at 10^6^: adsorption rate and growth rate. In those cases, close to 10^9^ phages are required at one extreme of the spectrum but only a few phage particles are enough at the other extreme. This means that a phage with a ‘good’ value of that phenotype is, on average, vastly more useful than a phage with a ‘bad’ value – that orders-of-magnitude fewer phage need be added to suppress the bacteria. Thus knowing the value of the phage property greatly informs the practitioner about whether billions or just a few phage will be needed.

In other cases, the variation in *P*_0_ across the X-axis is gradual. Here it may seem that measuring phage characteristics is of little value. Yet a difference of 1 in log(*P*_0_) is a 10-fold difference in the number of phage that must be administered; a log difference of 3 is a 1000-fold change in the number of phage needed. Therefore, even these gradual cases can have clinical importance.

A reasonable conjecture is that some of these properties interact – that knowing two phenotypes will be more predictive than knowing just one. Given the obvious utility of adsorption rate and the modest utility of decay rate, they seem a reasonable duo to consider. Fig. 3 is a contour plot of *P*_0_ per adsorption rate and decay rate (for cells at 10^6^). From the mostly vertical contours, it is easily seen that adsorption rate mostly determines the number of phage that must be added, but that there is a modest effect of decay rate. These are the conclusions that were inferred from considering those two phenotypes in isolation.

**Figure 3.**
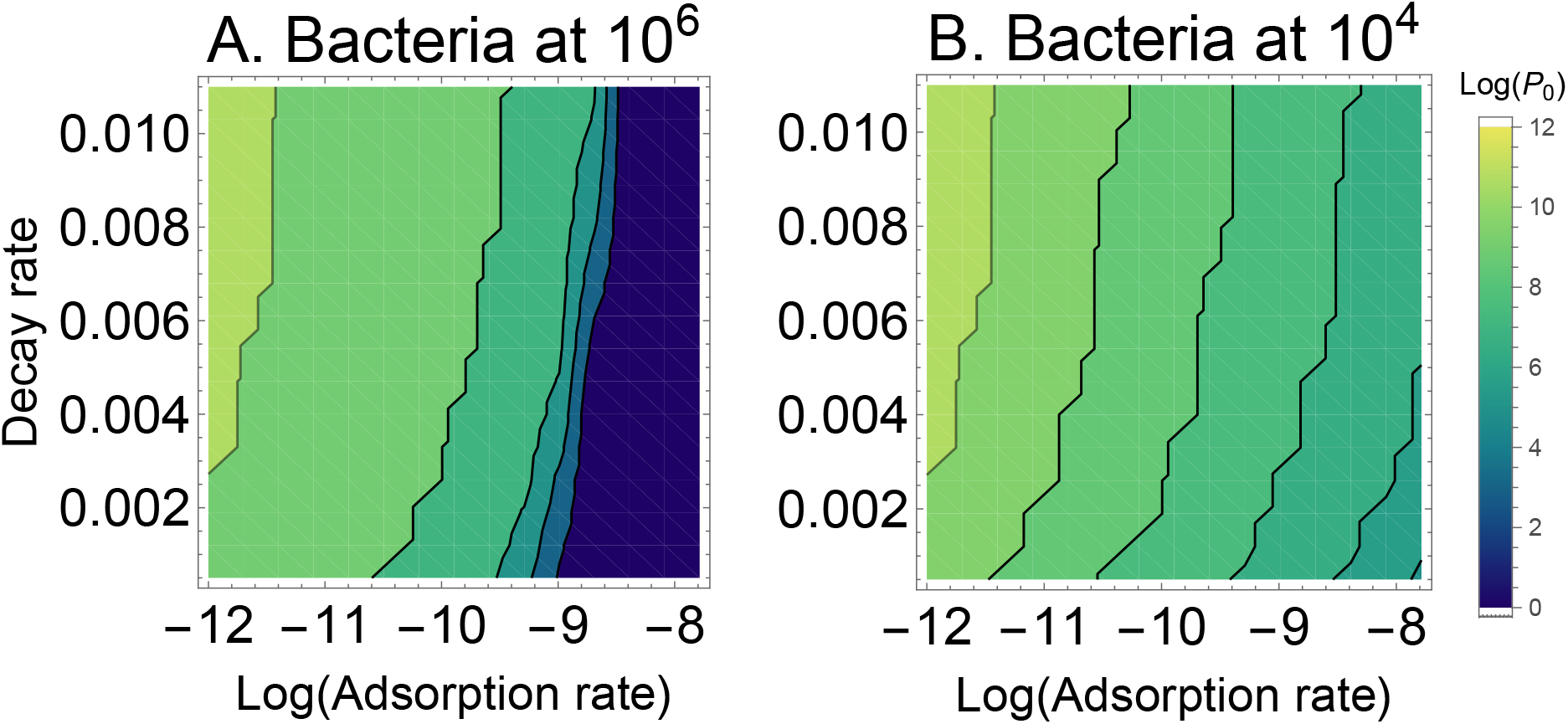
Heatmap of the log_10_ density of phage needed to suppress an initial bacterial density by 100-fold when varying adsorption rate (X axis, logged) and decay rate (Y axis). (A) Initial bacterial density is 10^6^. (B) Initial bacterial density is 10^4^. The contours span a vast range of 10^10^ to 1 in absolute numbers. Although there is the expected effect of bacterial density on the number of phage that need be added (*P*_0_), within a bacterial density, the difference in *P*_0_ is due mostly to adsorption rate. However, when considering the entire span of decay rate shown, there is an effect of 1-2 logs. These trials assumed a burst size of 20 and a lysis rate of 0.03.

For assessing the utility of decay rate, a more realistic model may be that of our eqn set (3), in which phage are added to a bacteria-free compartment from which they diffuse into the compartment with bacteria– as when phage are added to a patient at a site remote from the infection. The longer that the phage remain in the bacteria-free compartment, the more that decay is likely to affect phage success. However, even when assuming that all phage are added to the bacteria-free compartment and the migration rate out of the bacteria-free compartment is 0.001 (an average residence time of over 16 hours), decay rate becomes only modestly more important in phage success: at a cell density of 10^4^, the average *P*_0_ ranges from 8.3-9.9 (versus 7.8-8.8 in the single-compartment model), and at a cell density of 10^6^, the average *P*_0_ ranges from 6.7-8.7 (versus 6.0-7.6 in the single-compartment model). Thus there is indeed some increase in the utility of decay rate as a general measure of phage utility, but the increase is modest – and the compartment increases the necessary phage dose by several-fold. Adsorption rate and growth rate remain superior.

## Discussion

This study used computational models of bacteria-phage dynamics to assess the relative merits of different phage characteristics for use in phage therapy. The models assumed a nearly static bacterial population, indicative of a chronic infection, and numerically investigated the number of phage needed to suppress the bacterial population 100-fold inside of 1.4 days. The best predictors of phage success were adsorption rate and growth rate, the latter a composite measure of adsorption rate, lysis rate, and burst size that is nonetheless easily measured. The model included a phage decay rate with empirically-motivated parameter values, but decay rate was found to be less indicative of phage success than were adsorption and growth rates. To use these results, even qualitatively, one would need to characterize individual phages for adsorption rate and/or growth rate and apply the phages that had the best characteristics. It would be necessary to measure bacterial densities in the infection to know how much of a phage to add.

### Microplate models

An obvious comparison for our study is that of Blazanin et al. [3], who modeled phage growth in microplates, a high-throughput, practical method that would use microplate readers of optical density of cells (both uninfected and infected) as the time-dependent readout. The intent in that study was to know how to indirectly identify phages that may be good versus bad in therapy rather than to directly represent infections. The modeling served the same overall purpose as here – to discover which features of the phage growth and life history may reflect the ability of a phage to suppress the bacterial population.

Before comparing results, it is useful to consider some differences in the modeling approaches of the two studies. In their study [3], bacteria and phage were started at low densities, with bacteria growing to at least 10^8^ unless suppressed by phages. Eventual attainment of high bacterial density ensured that even a low density of a poor phage would eventually overwhelm the bacteria. In contrast, we assumed slow-growing bacteria whose density during treatment would be close to the initial density, hence rapid suppression of the bacteria might require addition of a very high density of phage. We likewise considered both a low and intermediate density of cells, to represent different kinds of infection. Second, there was no intrinsic phage death in their model (only loss to infection and superinfection), whereas our model allowed a decay/death rate. Our model thus accommodates known and possibly important mechanisms of in vivo phage loss [30]. Third, and related to the first point, our study applied a treatment-motivated measure of the effort required to suppress the infection. Our output was the number of phage needed to suppress the infection (100-fold suppression in a ∼ 1.5 day interval), so one could directly compare the practicality of using phages of different quality.

Given the differences in the modeling approaches of these studies, it is encouraging that several results are in common. Thus, both studies observed that adsorption rate was highly predictive of phage quality. Likewise, phage growth rate by any of several measures was also highly predictive of phage quality. In contrast, burst size was predictive of phage quality in their study but not in ours; some of this difference could be due to the smaller dynamic range of burst size used here, but the difference may also stem from the low cell densities used here, as we observed a substantial increase in the predictive utility of burst size when going from an infection density of 10^4^ to 10^6^. Thus, the commonality of several results between these studies suggests that various types of models may be used to obtain insight to phage properties useful in therapy.

In addition, however, the comparison of both studies identifies matters that will benefit from further analysis. One is translating phage quality into treatment practices. We converted our results into a phage dose that would be required to achieve a rapid, threshold suppression of bacterial density. Any such translation will be somewhat arbitrary, but it has the benefit of providing a semi-quantitative measure of treatment effort, and there will be many contexts in which it is desirable to know the consequences of applying a ‘good’ versus ‘bad’ phage. For example, one of the measures analyzed by Blazanin et al. [3] was the peak bacterial density attained before phage growth drove the bacterial numbers down. One could trivially (and arbitrarily) assign a threshold density of, say, 10^8^ as the boundary between treatment success and failure. But it is then important to know what protocol modifications could be implemented to attain treatment success with a phage that let cell density exceed 10^8^ in a microplate culture – or what reductions in effort would be enabled with a phage whose peak density was 10^7^.

In the same vein, it will be important to know bacterial densities of infections. The comparison of Blazanin et al. [3] to our study reveals that (relative) phage quality measures are somewhat independent of bacterial densities – that what constitutes ‘better’ is robust to the different models. But our study revealed that the number of phage that need be added is profoundly dependent on bacterial density – if bacterial density is not increasing rapidly. Low bacterial densities are common in sepsis [22] and some urinary tract infections [36], and it would be a mistake to admininister phages to those infections and assume that phage will amplify themselves to overwhelm the bacteria. So the use of models to identify relative phage quality is a first step, and it will be important to extend the work to infer absolute phage quality.

### Strictly in vivo properties of phages

The standard phage attributes of burst size, adsorption rate and lysis time – and composites such as growth rate – will have outputs in vitro, but some phage properties will apply only in vivo. One purely in vivo phenomenon is phage decay rate – phage loss due to capture by the host’s reticulo-endothelial system and by adherence to eukaryotic cells [30]. If decay rate cannot be measured in vitro, is there a reason to include it in the models?

Decay rate has been suggested to be an important determinant of success [30, 37]. Even if measuring decay rate is impractical, and even if decay rate can be only approximately parameterized in the models, the models can give an indication of its importance to phage performance, and thus whether the effort to measure would be justified. The results here suggest that decay rate has some predictive value but that other phage phenotypes are more important.

Using mice, Merril et al. [30] specifically considered decay rate in the context of prophylactic phage treatment (which contrasts with our assumption that bacteria are present from the start). For phage retention after 1 or more days prior to challenge with bacteria, decay rate can be profoundly important because decay follows *e*^−1440*δ*^ per day (where *δ* is the per-minute decay rate). In addition, Merril et al. [30] showed that decay rate could be evolved with simple selection for persistence in mice. Serwer [37] proposed that decay rate depends on phage electrical charge, thus providing a possible easy in vitro assessment. So if large differences in decay rate are present among wild phages or if large differences are easily evolved in vitro, this would be a robust phenotype to apply in the choice of phages, and it will have some benefit. And the example highlights the use of models to identify the utility of a phage phenotype that is intractable with in vitro growth assays.

Other in vivo properties will be omitted from all but the most specific models. For example, capsular depolymerases encoded by phages may greatly augment treatment by assisting immune clearance of the bacteria rather than by facilitating phage killing [e.g., 41, 40, 31, 8, 10]. At this stage, it is not clear even how to model such effects, but the benefits of using phages with depolymerases have been so profound in experimental work that modeling may be unnecessary.

### Feasibility of characterizing or evolving phages

How practical is it to characterize or ‘type’ a library of phages for the properties of interest? Might phages even be evolved to improve a property that is deemed worthwhile? If a property can theoretically reveal the therapeutic utility of a phage, the property is useful for large libraries only if it can be easily used to characterize phages. Burst size, adsorption rate, lysis time and growth rate all have well established methods for estimation [2], albeit that typing an entire library of 100 or more phages by standard methods would be tedious. Recent work suggests that any of various measures obtained from microplate readings (and other high-throughput methods) may suffice to characterize phage growth rates and that some individual phage phenotypes may be inferred from simple growth curves [28, 19, 3].

In vitro selection of higher adsorption should be practical, as phages are routinely selected for host range changes [e.g., Appelmans’s protocol, 29, 12]. Essentially any exposure of bacteria to phages in liquid should favor high adsorption, but growth in plaques does not [46]. However, selection for broad host range – for which Appelmans’s is typically applied and which offers the obvious benefit of breadth in treatment – may compromise adsorption to individual hosts by favoring breadth. The evolution of weak individual adsorption rates is a potential drawback to protocols favoring the evolution of broad host ranges. If adsorption to a particular host seems inadequate for a phage, selection could be applied for growth on just that host.

The possibility of using within-host evolution phages during treatment to acquire better phages for treatment has been considered using computational models [11]. Within-host evolution is moderately slow to improve therapy during the course of a single treatment. Furthermore, it requires repeatedly recovering phages from patients after treatment and then using those phages for continued or future treatment – avoiding repeated application of the same, initial stock. Multiple dosing with the same stock overrides any within-host phage evolution that would benefit treatment. So applying within-patient evolution to improve phages is moderately impractical.

### Translating model results into practice

There is no assurance that a phage’s ability to kill cells in a microplate or flask will reflect its ability to cure an infection. It is certainly encouraging that successful phage therapy often requires knowing little more than that a phage plaques on the bacterium of interest. But the issue raised here and by recent studies is whether differences in phage quality for treating an infection are correlated with phage ‘quality’ as measured in microplates or other in vitro system. It could be that the phages most suitable for treatment are not necessarily the best growers in vitro. In a strict sense, this question can only be answered by comparing phage performance in vitro to phage performance in vivo. But, as a first step, a simpler task would be to compare phage performances across different in vitro systems, such as planktonic growth, biofiilms, and other forms of spatial structure [14, 24, 42, 38]. If the ranking of different phages depends on the in vitro growth environment, then it is likely to differ between in vitro and infection environments.

We imagine three steps in applying these results to actual phage therapy. The most immediate one is to measure phages for the properties that best reflect success in the model, which appears to be adsorption rate or growth rate. These measurements would perhaps be done most easily on a large scale with microplate readers to quantify phage growth rate [19, 3]; assays of adsorption rate do not easily lend themselves to high throughput, even though adsorption rate appears to be equally good a predictor as growth rate. The phages with the best growth rates, or other suitable measure, would be used for treatment.

The second step is to decide which type of infections are good fits for the models (e.g., sepsis, urinary tract infections, abscesses, infection of hard tissues, …). This step might well precede phage typing. This is the most challenging step, because it confronts the huge gap between phage growth in idealized environments and phage growth in an infection – and the translation of that growth into infection control. Bacterial infections are typically heterogeneous, with bacteria varying in density and physiological state throughout the body. Some bacterial states or loci may be recalcitrant to infection by phages, others not [43, 39, 23]. If phages differ in how they access and grow in different compartments of the infection, which seems likely, then simple measures such as growth in planktonic cultures may yield poor insight to infection control. This step is important yet potentially intractable given the current state of knowledge. Practicality may dictate that we do not attempt to figure out whether an infection type fits the model, that any infection is treatable with the ‘best’ phages.

The third step is choosing a dose to administer. The models here were used to derive doses for treatment. But any such calculation is highly sensitive to quantitative details – in contrast to the apparent robustness of different measures of phage growth in suppressing the bacteria. Thus, the results from our study may well provide qualitative insight to whether an infection will require passive or active therapy, and thus requires an inoculum of many or few phage. But we caution against using the results quantitatively. A survey of recent phage therapy cases suggests that treatments rarely if ever rely on active therapy, with doses of 10^10^ and higher being common [15]. Thus, the dose administered may commonly be the highest dose that is practical given the constraints of phage preparations.

The many unknowns associated with applying simple models of phage performance to treatment are likely to be overcome only with data from actual treatments. It should be possible to monitor bacterial and phage densities before and after treatment, which would minimally inform whether the treatment falls into the realm of active versus passive therapy. Yet to actually compare treatment outcomes between phages with different model-based rankings is practical only with experimental infections. Indeed, phage success in vivo has not always aligned with in vitro growth in experimental, acute infections [e.g., 10, 21]. Thus it is already clear that experimental studies will be essential in evaluating whether and when the models have therapeutic utility. But it should also be remembered that the models – when appropriately crafted – can provide insight that will be difficult to attain empirically and can expedite treatment choices and help develop intuition. Thus, despite the many unknowns at the empirical level, the modeling has a role in improving phage therapy.

## Acknowledgments

We thank Steve Abedon for advice on the literature. GK was supported by an REEU grant from the University of Idaho, JB by P20-GM104420 from the National Institutes of Health.

